# A549 tumorigenic and BEAS-2B non-tumorigenic cell line derived small extracellular vesicles show distinct proteomic, *N*-glycoproteomic and chondroitin/dermatan sulfate profiles

**DOI:** 10.1101/2025.03.13.643059

**Authors:** Mirjam Balbisi, Tamás Langó, Virág Nikolett Horváth, Domonkos Pál, Gitta Schlosser, Gábor Kecskeméti, Zoltán Szabó, Kinga Ilyés, Nikolett Nagy, Otília Tóth, Tamás Visnovitz, Zoltán Varga, Beáta G. Vértessy, Lilla Turiák

## Abstract

Extracellular vesicles (EVs) are critical mediators of intercellular communication and hold promise as biomarkers and therapeutic targets in cancer, but their molecular alterations remain poorly understood. Protein glycosylation is a frequent post-translational modification; however, most EV studies focus only on proteomics, while mapping glycosylation changes of proteins are still underrepresented. To address this shortcoming, we analyzed the proteomic, *N*-glycoproteomic, and chondroitin/dermatan sulfate (CS/DS) glycosaminoglycan (GAG) profiles of small EVs (sEVs) derived from A549 lung adenocarcinoma and BEAS-2B non-tumorigenic epithelial cell lines. Principal component analysis and hierarchical clustering revealed that all three profiles are highly dependent on the origin of sEV, highlighting fundamental differences not only at the proteomic but also at the *N*-glycopeptide and CS/DS levels. Protein expression differences were primarily associated with the upregulation of cell cycle regulation, DNA repair, metabolism, and protein synthesis, while immune-related processes were predominantly downregulated. Proteomics revealed differential expressions of 5 CS proteoglycans, anticipating that their CS profile may also change. *N*-glycoproteomics highlighted a shift from complex to hybrid *N*-glycans in cancer sEVs, alongside a significant decrease in fucosylation. Prominent glycoproteins characterized with multiple glycosylation sites included versican, galectin-3-binding protein and laminins. The total amount of CS/DS increased 3.4-fold in cancer sEVs, while the ratio of the two monosulfated disaccharides changed 2-fold, suggesting altered sulfation mechanisms. These findings highlight the potential of *N*-glycoproteomics and GAG profiling to enhance biomarker discovery and EV-based cancer diagnostics.

**Graphical abstract:** 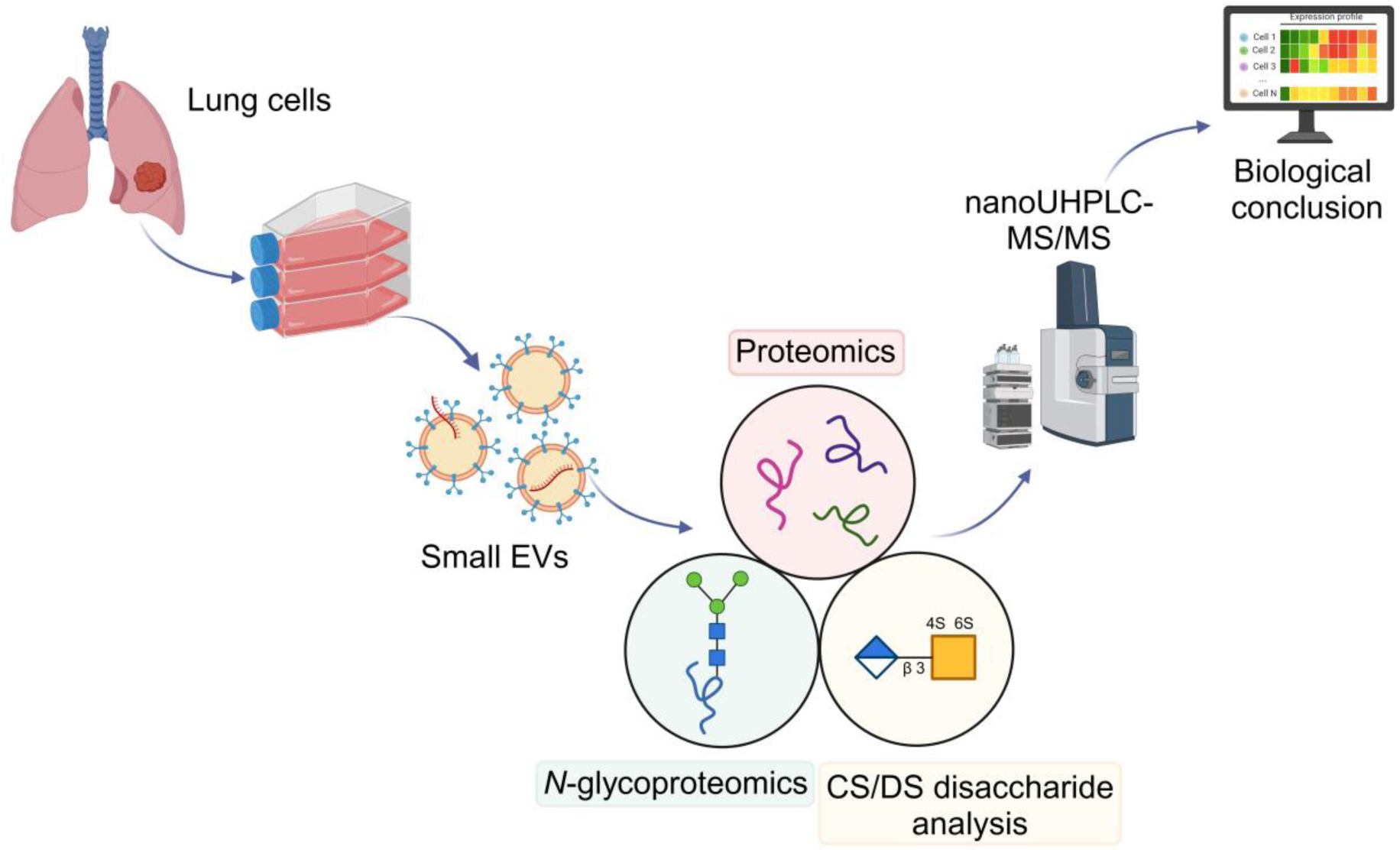

Proteomic, *N*-glycoproteomic and chondroitin/dermatan sulfate disaccharide profiles differ between A549 lung adenocarcinoma and BEAS-2B non-tumorigenic epithelial cell derived small extracellular vesicles.

## 1. Introduction

Lung cancer is a severe disease accounting for almost 2.5 million cases and 1.8 million deaths each year worldwide[1]. One reason for the high mortality rate is that the diagnosis is often made at an advanced stage of the disease, when surgery is no longer possible[2]. Another major factor is that although there are now several therapeutic options available for patients with certain genetic[3] or immunological[4] features, the choice of therapy is not straightforward and often requires tissue biopsies to identify specific/targetable genetic alterations. Therefore, there is an urgent need to develop less invasive methods to detect lung cancer and identify features for therapy selection, e.g. based on extracellular vesicles (EVs) in blood and other body fluids[5, 6].

EVs are lipid bound particles released from cells into the extracellular space that play a significant role in intercellular communication and carry various biomolecules, including proteins, lipids, and nucleic acids[7, 8], influencing the tumor microenvironment, cancer progression and metastasis[9, 10]. Based on their size, EVs can be classified into small (< 200 nm) and large EVs (> 200 nm), of which small EVs (sEVs) are more commonly studied. Biologically, sEVs modulate several processes during tumor development, such as angiogenesis, cell transformation, invasion, metastasis, immune escape, and drug resistance[11, 12]. Tumor-derived sEVs can be detected in various body fluids, such as blood and urine, making them not only potential targets for future cancer treatments, but also a source of potential biomarkers for cancer detection and progression monitoring[13]. The primary molecular targets for this are nucleic acids and proteins, while specific modifications on EV proteins remain largely uncharacterized.

Over the past three decades, MS-based bottom-up workflows have revolutionized proteomics by enabling the simultaneous identification and quantification of large numbers of proteins[14, 15]. Proteomic profiles of EVs are widely studied in several types of cancer, e.g. breast[16], colorectal[17], prostate[18] and lung cancer[19]. It has also been shown that tumor-derived plasma EV proteomics has the potential to discriminate between different cancer types[20]. Proteins carry several post-translational modifications (PTMs) affecting their structure and function. However, the PTMs occurring on EV proteins are still relatively underexplored[21, 22]. Protein glycosylation is observed in over 50% of human proteins and is integral to several biological processes, influencing both cellular interactions (cell-cell interaction, signal transduction, immune response) and protein dynamics (protein folding, molecular recognition)[23].

Major protein glycosylation subtypes include *N*-glycosylation and *O*-glycosylation. A special class of glycosylated proteins are the proteoglycans (PGs), in which glycosaminoglycans (GAGs) are attached to a core protein. In the current study, *N*-glycosylation and chondroitin/dermatan sulfates (a class of GAGs) are investigated.

Human *N*-glycans have a common pentasaccharide core structure of 2 *N*-acetyl-glucosamine (GlcNAc) and 3 Mannose (Man), which can be extended into high mannose, complex and hybrid types. The *N*-glycoproteome can be analyzed either by enzymatic cleavage of the glycan chain, resulting in an average pattern (*N*-glycomics), or by enzymatic cleavage of the proteins and enrichment of the glycopeptides from the peptide mixture, resulting in site-specific information (*N*-glycoproteomics)[24]. Aberrant *N*-glycosylation was observed in several cancer types, including lung[25], colorectal[26] and breast[27] cancer. Several tumor biomarkers approved for clinical use are also glycoproteins, e.g. alpha-fetoprotein for liver cancer, prostate-specific antigen for prostate cancer, and carcinoembryonic antigen for colorectal cancer[28]. GAGs are long, linear polysaccharides of repeating disaccharide units that are divided into classes based on the structure of the disaccharides[29]. Chondroitin/dermatan sulfate (CS/DS) consists of glucuronic acid/iduronic acid and *N*-acetylgalactosamine units, and the disaccharides can be sulfated at C4 and C6 positions of GalNAc and less frequently at C2 position of uronic acid[30]. Chondroitin/dermatan sulfates are typically analyzed by bottom-up techniques, which involve bacterial lyase enzymatic digestion of the chains followed by high pressure liquid chromatography-mass spectrometry (HPLC-MS) to identify and quantify the different disaccharides present in the sample[31]. GAGs are present in different amounts and have different rates of sulfation compared to healthy controls in various cancer types, including lung[32], prostate[33] and liver[34] cancer.

The glycan composition of EV proteins can significantly influence the functional efficacy of EVs and altered glycan profiles can play an important role in cancer development. In the present study, we characterized the proteomics, *N*-glycoproteomics and CS/DS GAG profiles of sEVs derived from A549 lung adenocarcinoma cell line representing the most common lung cancer subtype[35], and BEAS-2B epithelial cell line isolated from non-cancerous bronchial epithelium. Exploring the glycosylation differences between tumor and non-tumor EVs may help to understand the role of *N*-glycosylation and CS/DS in cancer pathogenesis and provide further insights into the underlying biological processes.

## 2. Materials and methods

### 2.1. Materials

A549 and BEAS-2B cells, Trypsin-EDTA solution, phosphate buffered saline (PBS), poly-L-lysine (PLL), ammonium acetate, ammonium bicarbonate (AmBic), ammonium formate, formic acid (FA), trifluoroacetic acid (TFA), iodoacetamide (IAA), dithiothreitol (DTT), and chondroitinase ABC were obtained from Merck (Darmstadt, Germany); LC-MS grade acetonitrile, water, and methanol (MeOH) were purchased from VWR (Radnor, Pennsylvania, USA). Pierce C_18_ and graphite spin columns, Ham’s F-12 Nutrient Mix (supplemented with GlutaMAX), fetal bovine serum (FBS) and penicillin-streptomycin (PS) were acquired from Thermo Fisher Scientific (Waltham, Massachusetts, USA). Trypsin/Lys-C and Trypsin Gold were obtained from Promega (Madison, Wisconsin, USA). Econo-Pac chromatography columns were purchased from Bio-Rad (Hercules, California, USA), and Sepharose CL-2B from Cytiva (Marlborough, Massachusetts, USA). Rapigest SF Surfactant was acquired from Waters Corporation (Milford, Massachusetts, USA), CS disaccharide standards from Iduron (Manchester, UK) and bronchial epithelial growth medium (BEGM) BulletKit from Biocenter (Szeged, Hungary). Acetone was purchased from Honeywell (Charlotte, North Carolina, USA), and ammonia (25%) from Reanal (Budapest, Hungary).

### 2.2. Cell culturing

A549 cells were grown and maintained in F12 medium completed with 10% FBS and 1% PS inside a humidified incubator with 5% CO_2_ at 37°C (Eppendorf, Galaxy 170R). BEAS-2B cells were maintained in BEGM medium in similar conditions, after the flasks were pre-coated with 0.01% PLL for 20 min and washed with water twice. 72 hours before the start of sEV isolation, to minimize FBS contamination, 10-10 ml of serum-free medium (non-completed F12 and completed BEGM) was added to two 50% confluent T75 flasks to obtain one sample. Passage numbers of the cells used for sEV isolation are shown in Supplementary Table S-1. To avoid misidentifications due to differences in cell culture media, F12 and BEGM media samples were also prepared as control for proteomic, *N*-glycoproteomic and GAG analyses. 6-6 sEV biological replicates and 3-3 control media samples per cell type were analyzed.

### 2.3. sEV isolation

After 72 h of incubation, cell culture supernatants were collected and sEVs were isolated as described previously[36], with slight modifications. Briefly, solutions were centrifuged at 10000 g, 4 °C for 30 min to remove cell debris, apoptotic bodies and microvesicles, and supernatants were filtered through a 0.22 μm filter. The filtrates were concentrated to 1 ml on 10 kDa Amicon Ultra-15 centrifugal filters at 5000 g, 4 °C and sEVs were isolated from the resulting samples on in-house prepared size-exclusion chromatography (SEC) columns[36] filled with 10 ml Sepharose CL-2B, using 1 ml PBS to elute the fractions. Six fractions were collected, and based on preliminary size distribution and protein concentration measurements, fractions 3, 4 and 5 were combined for further analysis.

### 2.4. sEV characterization

#### Microfluidic resistive pulse sensing measurements (MRPS)

Fractions 3, 4 and 5 containing sEVs were combined and the mixture was diluted 5-fold with PBS containing 0.3 w/v% Tween-20, filtered through a 100 kDa Vivaspin 500 membrane filter. Samples with a volume of 5 μL were pipetted into C400 cartridges and measured using an nCS1 instrument (Spectradyne LLC, Signal Hill, California, USA) with a measurement range of 65-400 nm.

#### Transmission electron microscopy (TEM)

Samples for TEM were prepared as described previously[37], with slight modifications. Briefly, 3 μL of samples were deposited on formvar-coated grids, and dried for 10 min. EVs were fixed with 2% glutaraldehyde in PBS for 10 min and washed three times with water for 5-5 min. EVs were contrasted with 2% methyl cellulose containing 0.4% UranyLess (Electron Microscopy Sciences, Hatfield, Pennsylvania, USA) for 10 min on ice. Measurements were performed on a JEM1010 (Jeol, Japan) transmission electron microscope and images were analyzed by ImageJ. Diameters of sEVs (*N* = 50-50) were determined.

### 2.5. Solvent exchange and lysis

Collected SEC fractions were concentrated, and the solvent was exchanged to MS compatible AmBic buffer. Therefore, 10 kDa Amicon Ultra-0.5 centrifuge filters were first washed with 200 μL of water and centrifuged at 13500 g for 10 min. The sample was then pipetted onto the filters in 500 μL units and centrifuged at 13500 g for 10 min in each step. Subsequently, 200 μL 200 mM AmBic solution was added and centrifuged at 13500 g for 10 min, followed by the addition of 200 μL 50 mM AmBic solution and centrifugation at 13500 g for 15 min. Finally, the filters were turned upside down and centrifuged at 1000 g for 1 min to collect the sEV fractions. The sEVs were lysed with 7 consecutive freeze-thaw cycles, using 30 seconds of liquid nitrogen (cycles 1, 3, 5, 6, 7) or 1 hour of freezing (cycles 2, 4), followed by 10 min of ultrasonication each time. Protein concentrations were measured on a NanoDrop ND-1000 instrument (Thermo Fisher Scientific, Waltham, Massachusetts, USA) at 280 nm using bovine serum albumin calibration solutions from 0.1 μg/μL to 10 μg/μL. In cases where protein amounts were <15 μg (commonly observed for BEAS-2B sEVs), two samples were combined for further analysis. Further sample preparation steps were performed on 6-6 sEV samples of both cell types and 3-3 control media samples.

### 2.6. Proteomics and *N*-glycoproteomics workflow

#### Proteomics digestion

5 μg protein amounts of each sample were diluted to 35 μL with water, 1.5 μL MeOH, 2 μL 200 mM DTT and 5 μL 0.5% Rapigest were added and incubated at 60 °C for 30 min. 2.5 μL 200 mM IAA and 5 μL 200 mM AmBic solution were added to the samples and incubated at room temperature in the dark for 30 min. The digestion was performed in two consecutive steps: first, 50 ng of trypsin/Lys-C mixture was added to the samples and incubated for 1 h at 37 °C, followed by incubation with 500 ng of trypsin enzyme for another 15 h at 37 °C. Finally, the digestion was stopped by adding 0.5 μL FA and the peptide samples were dried down.

#### *N*-glycopeptide enrichment and peptide purification

For enrichment of *N*-glycopeptides, acetone precipitation was used, by first dissolving the peptide samples in 15 μL 1% FA, then adding 150 μL ice-cold acetone, and finally storing the samples in the freezer for 18 h[38]. The samples were then centrifuged at 12000 g for 10 min at 20 °C, and the supernatants (mostly non-glycosylated peptides) were separated from the pellets (mostly glycosylated peptides). Both fractions were dried down and the non-glycosylated peptides were purified on C_18_ spin cartridge. In short, the cartridge was washed with 2 × 200 µL 50% MeOH, 2 × 200 µL 0.5% TFA + 5% ACN, and 2 × 200 µL 0.1% TFA solution. Samples were applied in 50 µL 0.1% TFA solution and reapplied once. Contaminants were washed away with 2 × 100 µL 0.1% TFA, and peptides were eluted with 2 × 50 µL 0.1% TFA + 70% ACN solution. All steps were performed at 2000 rpm for 2 min. Elution solvents were evaporated, and both peptides and *N*-glycopeptides were stored at −20 °C until further use.

#### nanoUHPLC-MS/MS measurements for proteomics

Proteomic samples were dissolved in 0.1% FA + 2% ACN solution, and 200 ng was injected from each sample. Samples were analyzed on a timsTOF HT (Bruker Daltonics, Bremen, Germany) coupled with a Dionex Ultimate 3000 nanoUHPLC (ThermoFisher Scientific, Waltham, Massachusetts, USA). Samples were first loaded onto an Acclaim PepMap C_18_ trap column (5.0 μm, 300 μm × 5 mm, Thermo Fisher Scientific, Waltham, Massachusetts, USA) at a flow rate of 10 μL/min, followed by separation on a monolithic capillary MOSAIC C_18_ analytical column (75 μm × 150 mm, Bruker, Bremen, Germany) heated at 50 °C. Eluent A consisted of 0.1% FA in water, while eluent B was 0.1% FA in 80% ACN. The gradient started at 5% B and increased to 40% B over 20 min at a flow rate of 0.5 μL/min.

The MS ion source was a CaptiveSpray 1 source with a 10 µm emitter used in positive mode, with a capillary voltage of 1500 V, a mass range of *m*/*z* 100-1700 and an ion mobility range of 0.7-1.4 V·s/cm². To optimize data acquisition, data-dependent analysis (parallel accumulation-serial fragmentation, PASEF) was employed first. A spectral library was generated using FragPipe, data-independent acquisition (DIA) window optimization was carried out with py_diAID[39], and samples were measured by dia-PASEF. Transfer time was 60 µs and pre-pulse storage time was 12 µs. The TIMS settings included a ramp time of 180 ms and an accumulation time of 180 ms; PASEF was performed with 4 MS/MS scans per cycle and a total cycle time of 0.93 s. Precursors were selected within a charge range of 0 to 5, with an intensity threshold of 1500 for scheduling and a target intensity of 15000. Exclusion release time was 0.4 min, reconsider precursor switch was enabled and a current-to-previous intensity ratio of 4 was set. Exclusion windows were set to 0.015 *m*/*z* for mass width and 0.015 V·s/cm² for ion mobility width.

#### nanoUHPLC-MS/MS measurements for *N*-glycoproteomics

Glycoproteomic samples were dissolved in 10 μL 0.1% FA + 2% ACN solution, of which 2 μL was injected. Measurements were performed on a Waters nanoAcquity nanoUHPLC system (Milford, Massachusetts, USA) coupled to a Thermo Fisher Exploris 240 Orbitrap MS (Waltham, Massachusetts, USA). Chromatographic separation utilized a Symmetry C_18_ trap column (5 μm, 180 μm × 20 mm, Waters, Milford, Massachusetts, USA) and an Acquity M-Class BEH130 C_18_ analytical column (1.7 μm, 75 μm × 250 mm, Waters, Budapest, Hungary). Eluent A consisted of 0.1% FA in water, while eluent B was 0.1% FA in ACN, and flow rate was 300 nL/min. The gradient program started from 2% B (0-1 min), increased from 2% to 25% B (2-82 min), then from 25% to 40% B (82-85 min) and from 40% to 90% B (85-86 min), kept there for 2 min (86-88 min) and finally, the column was re-equilibrated at 2% B (88-90 min).

The MS was operated in positive mode with a capillary temperature of 275 °C and a capillary voltage of 1.8 kV. MS full scans were acquired at a resolution of 120000 in the mass range of 360-2200 Da, with an automatic gain control (AGC) target of 2×10⁶ to maintain consistent signal intensity and a maximum injection time of 200 ms. Ions were selected within a 2 Da isolation window to MS/MS, and stepwise higher energy collisional dissociation fragmentation energies of 10, 20, and 30 eV were used. The resolution was maintained at 120000, with an AGC target of 2×10⁵, a maximum injection time of 200 ms, and a mass range of 200-2000 Da. The minimum precursor intensity threshold was set to 1.7×10⁴ and the minimum AGC target to 10³.

#### Proteomics data evaluation and visualization

DIA-NN[40] was used to identify and quantify proteins, using the Uniprot human database (access date: 03.2024, 20434 sequences) and trypsin/P enzyme. Carbamidomethylation of cysteine amino acids was set as a fixed modification, while methionine oxidation, *N*-terminal methionine excision and protein *N*-terminal acetylation were set as variable modifications. A maximum of 1 missed cleavage site and 1 variable modification was allowed. Detailed settings are shown in Supplementary Table S-2.

Statistical evaluation and visualization of the results were performed with custom code in R[41] 4.3.2 using RStudio[42] 2024.12.1+467. Proteins identified with less than 2 unique peptides or detected in at least 1/3 of the control media samples were excluded and only those proteins quantified in at least half of the samples in at least one sample group were considered for further analysis. Imputation of missing values was performed based on the number of detections in each group: if the given protein was detected in less than 2/3 of the samples in the group, sample’s 5-percentile was imputed, whereas in case of less missing values, it was imputed using the kNN algorithm (VIM package[43], k = 15). Normality and equality of variances were tested on log-transformed data using Shapiro-Wilk and Levene tests, respectively and based on the outcome, Student’s *t*-test, Welch *t*-test, or Wilcoxon rank sum test was performed for the given protein. False discovery rates were controlled with the Benjamini-Hochberg method and adjusted *p*-values less than 0.05 were considered significant.

For visualization, the packages ggplot2[44] and gplots[45] were used, in which principal component analysis (PCA, prcomp function), hierarchical clustering (heatmap.2 function, ward.D2 method), volcano diagram and boxplots were generated. Gene set enrichment analysis (GSEA) was conducted using clusterProfiler[46] on ranked genes from statistical analysis, identifying enriched Gene Ontology (GO) Biological Processes (adjusted *p*-value cutoff = 0.1 was used).

#### *N*-glycoproteomic data evaluation and visualization

Glycoproteomics data were first searched in Byonic 5.0.20[47] against the Uniprot human database (access date: 03.2024, 20434 sequences), with MS1 mass accuracy of 7 ppm, MS2 mass accuracy of 20 ppm and 1% false discovery rate. Carbamidomethylation of cysteine was a fixed modification, whereas deamidation of asparagine and glutamine, oxidation of methionine and presence of *N*-glycans (Byonic’s built-in database, 182 human *N*-glycans without multiple fucose) were variable modifications. The number of missed cleavages was limited to 2, the maximum number of common modifications to 2, and the maximum number of rare modifications to 1. Glycopeptide hits were filtered for LogProb >2 and score >200, corresponding proteins were manually checked for known glycosylation sites, and only proteins confirmed to be *N*-glycosylated were used for further analysis. Next, *N*-glycopeptides were quantified using GlycReSoft[48] 0.4.22 with the protein list generated using the Byonic search and an in-house *N*-glycan database (160 glycans in total, no multiple fucose, see Supplementary Table S-3). Detailed settings of GlycReSoft searches are shown in Supplementary Table S-2. Results were imported into R and processed. *N*-glycopeptide assignments with MS1 score > 3 and MS2 score > 5 were accepted[49], cell culture media derived glycopeptides were removed as in the case of proteomics and intensities were normalized using total area normalization. Statistics and visualization of glycoproteomics data were performed in the same way as described for proteomics, and glycoforms were screened to ensure that they belong to known glycosylation sites. Glycosylation metrics were used to characterize sialylation (the ratio of sialylated antennae), galactosylation (the ratio of galactosylated antennae), fucosylation (the ratio of fucosylated glycopeptides) and the ratio of different types of glycopeptides.

### 2.7. Chondroitin/Dermatan sulfate workflow

#### CS/DS digestion

10 μg proteins were made up to 70 μL with water, to which 20 μL 500 mM AmBic solution, 5 μL 100 mM ammonium acetate and 5 μL 5 mU/μL chondroitinase ABC enzyme were added and incubated at 37 °C for 16 h. Digestion was stopped by placing the samples at 90 °C for 3 min and the samples were dried down and stored at −20 °C until the purification.

#### CS/DS disaccharide purification

For the purification of CS/DS disaccharides, we used a cotton-hydrophilic interaction chromatography (cotton-HILIC) + graphite solid-phase extraction two-step procedure developed in our group. In each step, samples were centrifuged at 2500 rpm for 1 min. In the first part, self-packed cotton-HILIC pipette tips were used, that were first washed with 50 µL 60% ACN solution, followed by 2 × 50 µL 1% TFA + 95% ACN solution. Samples were applied and reapplied twice in 30 µL 1% TFA + 95% ACN solution, then washed with 50 µL 1% TFA + 95% ACN, and eluted with 2 × 10 µL 1% ammonia solution pre-heated to 40 °C. Flow-through (from sample application and washing) and elution fractions were dried down, and flow-throughs were further purified on Thermo Pierce graphite cartridges. To do this, 2 × 100 µL 0.1% TFA + 80% ACN solution was used, followed by 2 × 100 µL water. Samples were applied and reapplied once in 50 µL water, washed with 3 × 100 µL water and eluted with 3 × 50 µL 0.05% TFA + 40% ACN solution. The elution fractions were combined with the cotton-HILIC elution fractions, dried down and stored at −20 °C until further use.

#### UHPLC-MS/MS measurements

CS/DS disaccharide samples were dissolved in 8 μL 10 mM ammonium formate + 75% ACN (pH 4.4) solution, of which 2 μL was injected. Samples were measured on a Waters Select Series Cyclic IMS (Milford, Massachusetts, USA) coupled to a Waters Acquity I-Class UPLC (Milford, Massachusetts, USA) equipped with a self-packed GlycanPac AXH-1 HILIC-weak anion exchange capillary column (250 μm × 10 cm). A and B solvents were 10 mM and 65 mM ammonium formate + 75% ACN (pH 4.4) solution[50]. The flow rate was 10 μL/min, and CS/DS disaccharides were separated with constant 5% B for 7 min, followed by washing with 95% B for 5 min and equilibration with 5% B for 3 min. Extracted ion chromatograms of characteristic ions for CS/DS disaccharides are shown in Supplementary Fig. S-1.

A low-flow electrospray source was operated in negative mode, with a capillary voltage of 2.5 kV, a cone voltage of 10 eV, and an ion source temperature of 120 °C. MS1 spectra were collected with a trap collision energy of 6 eV and transfer collision energy of 4 eV in the *m*/*z* 200-600 mass range, while MS/MS spectra were taken at 20 eV collision energy to differentiate between D0a4 and D0a6.

#### CS/DS data evaluation and visualization

TargetLynx integrated in MassLynx V4.2 software was used to integrate the chromatographic peaks of GAG disaccharides, and then curves were fitted in Microsoft Excel to calibration samples containing known amounts of CS disaccharides and the area of the biological samples was converted to fmol values. CS/DS disaccharides could not be detected in one A549 sample, presumably due to sample preparation errors, the sample was therefore excluded from the analysis. Results were plotted and statistically evaluated in R 4.3.2 using RStudio 2024.12.1+467 with custom code, similar to proteomics and *N*-glycoproteomics. In short, boxplots, PCA and heatmaps were used for visualization, and Student’s *t*-test, Welch *t*-test, or Wilcoxon rank sum test were used for statistics, and Benjamini-Hochberg correction was applied.

## 3. Results

6-6 parallel sEV samples were isolated from the cell culture supernatant of A549 adenocarcinoma and BEAS-2B non-tumorigenic epithelial cells, while cell culture media were used as controls. Samples were characterized according to the MISEV2023 guidelines[51]. An overview of the workflow is presented in Fig. 1. First, proteomic digestion was performed, and the resulting peptide mixture was enriched for *N*-glycopeptides, while the remaining fraction was used for proteomic analysis. In addition to MRPS and TEM analysis, DIA proteomics was used to verify the purity of sEVs by the presence of EV marker proteins. We then assessed protein expression differences between normal and tumor sample groups, with a special focus on PG core proteins. PCA and heatmap clustering were used to detect system-level differences, and GSEA was performed to identify altered biological processes. *N*-glycoproteomics results were interpreted using statistics, volcano plot, PCA and heatmap. In addition, we determined the distribution of different types of glycans and calculated glycosylation metrics to characterize overall *N*-glycomics changes. Glycoproteomic results were also correlated with proteomics. On the other hand, CS/DS GAG chains were digested into CS/DS disaccharides. The GAG disaccharides investigated in this study are shown in Fig. 2. The total amount of CS/DS disaccharides, their relative proportion, the D0a6/D0a4 ratio (hereafter: 6S/4S) and the average rate of sulfation were calculated, and PCA and heatmap analysis were performed.

**Figure 1.**
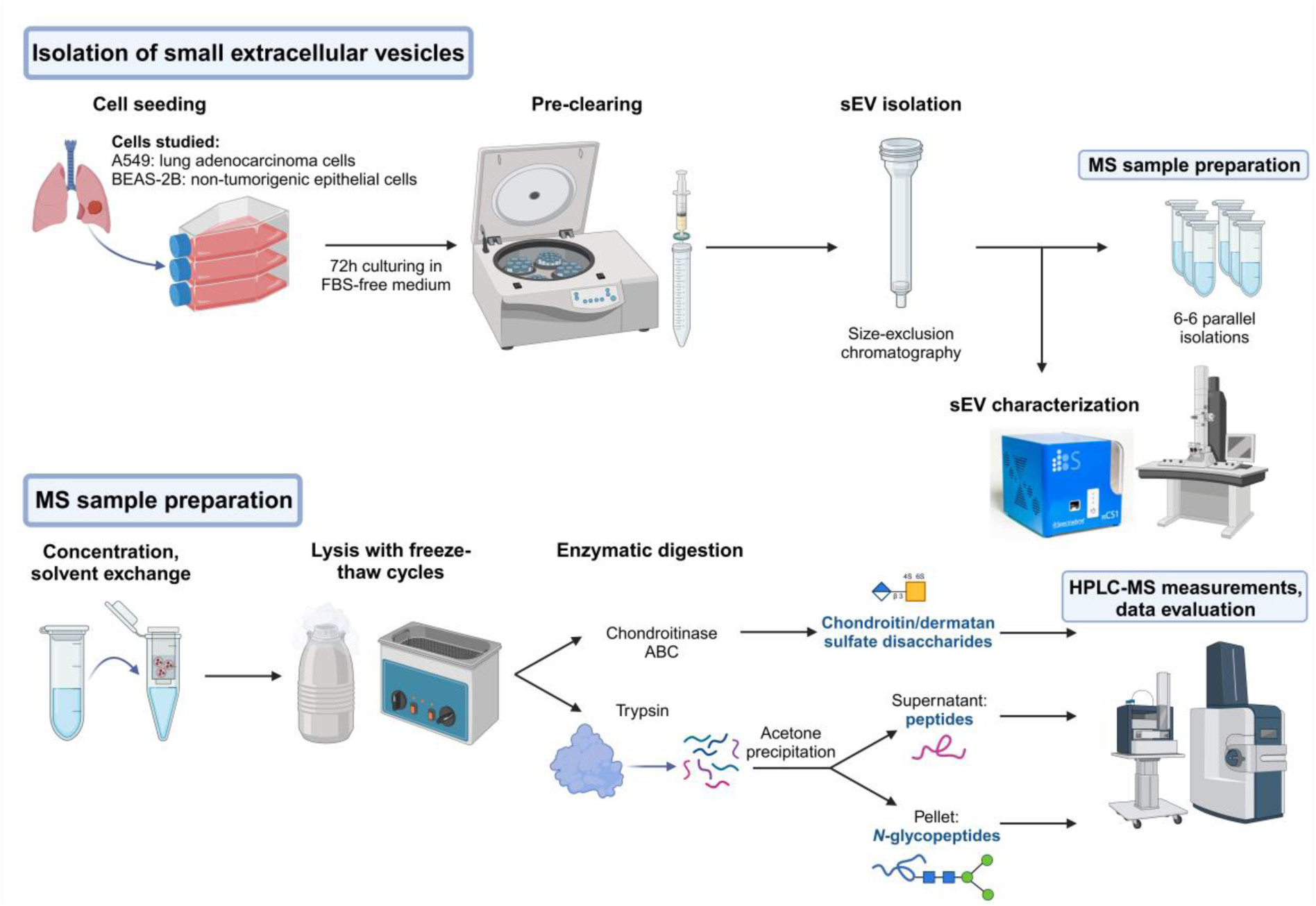
Workflow of sEV isolation and proteomic, *N*-glycoproteomic and CS/DS disaccharide analysis of A549 and BEAS-2B derived sEVs. Created with BioRender.com.

**Figure 2.**
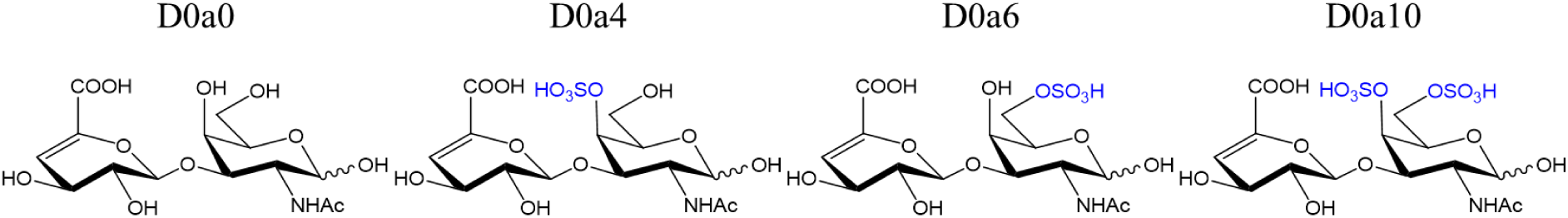
Lawrence codes and structures of CS/DS disaccharides produced during enzymatic digestion.

### 3.1. sEV characterization

First, we examined the size and shape of the isolated sEVs by TEM and characterized their size distribution by MRPS. Results for A549 and BEAS-2B sEVs are shown in Fig. 3/A and B, respectively. TEM analysis confirmed the presence of spherical particles with smaller than 200 nm size. In general, smaller particles were found in A549 samples than in BEAS-2B. In A549, most particles were between 30-120 nm in size according to TEM, while in BEAS-2B they were mainly between 50-250 nm (Fig. 3/C). MRPS analysis indicated slightly lower particle diameters, but the same trend in the size of the isolated particles, i.e. a higher proportion of larger particles was found in BEAS-2B sEVs than in A549 sEVs. Another difference was observed in the particle concentration, as A549 samples had a considerably higher number of sEVs. To reduce differences, enzymatic digestion was performed on equal amounts of protein.

**Figure 3.**
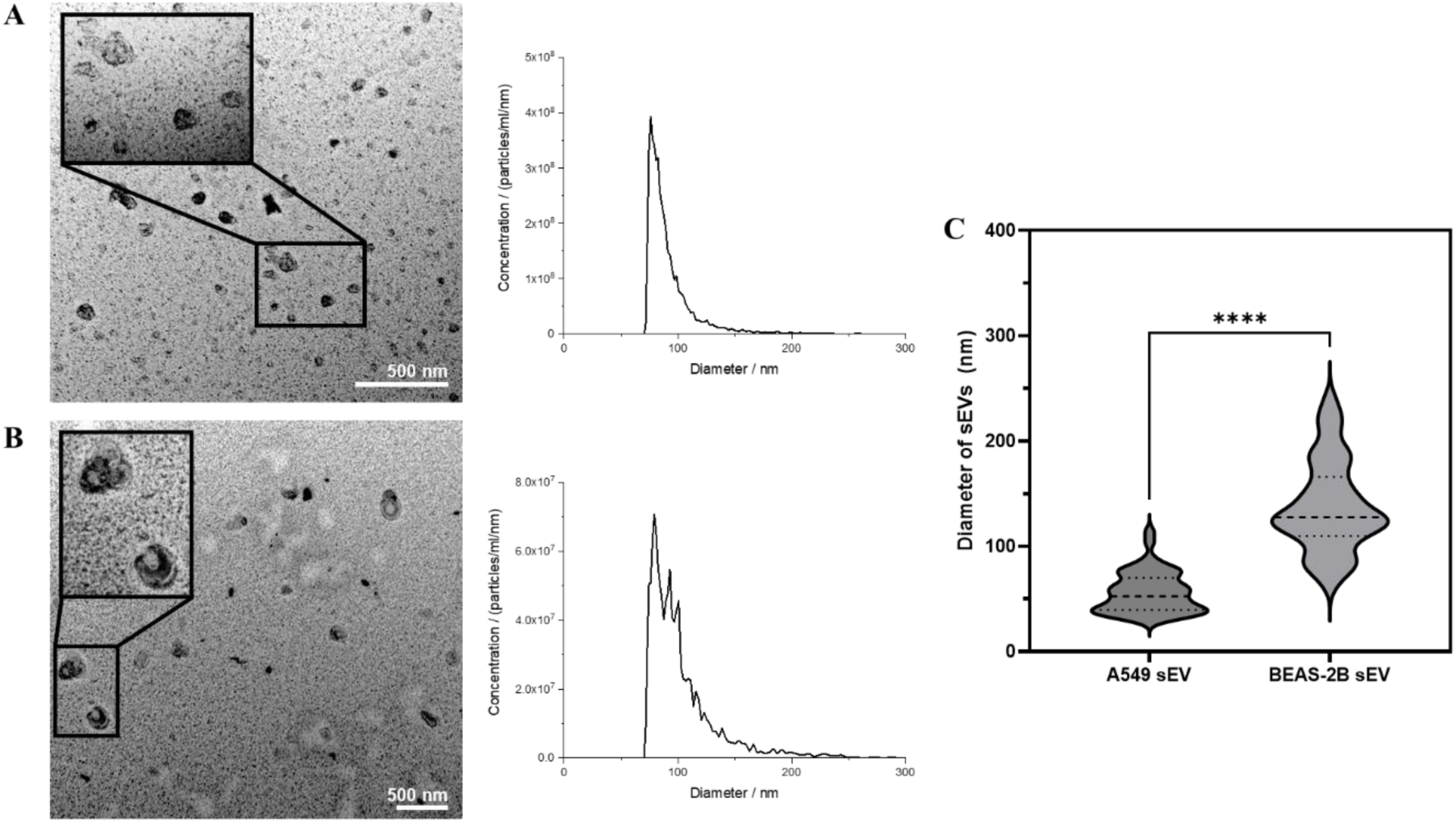
Transmission electron microscopy and microfluidic resisitive pulse sensing analysis of **A.** A549 derived sEVs. **B.** BEAS-2B derived sEVs. For MRPS, the average of the particle concentration of 3-3 samples is visualized. **C.** Distribution of the diameters of sEVs measured by TEM images (*N* = 50-50). (****p < 0.0001)

### 3.2. Proteomics

In the DIA label-free proteomic experiments, a total of 2334 proteins were identified and quantified from 12 sEV (6 A549 and 6 BEAS-2B) and 6 (3 F12 and 3 BEGM) media samples. Quantified proteins were compared to the 100 most frequently identified exosome proteins listed in ExoCarta[52], and each individual sample overlapped 77-91 hits from the list, demonstrating the good quality of sEVs isolated by SEC. 1528 proteins were detected in at least 3 replicates in at least 1 sEV group, but 583 of them were also present in at least 2 media samples causing the bias of observed expression levels in sEV samples. Therefore, only the remaining 945 proteins were included for statistical analysis, of which 408 were found to have different abundances between the two sample groups. Among them, 313 proteins were upregulated in A549, while 95 were downregulated. For example, top upregulated proteins were alpha-fetoprotein and aldehyde dehydrogenase 3 family member A, while top downregulated proteins were pentraxin-related protein PTX3 and fibulin-2. The distribution of fold-changes (FCs) and adjusted *p*-values is visualized on the volcano plot (Fig. 4/A). The full list of statistical results for the proteomic analysis can be found in Supplementary Table S-4. 11 PG core proteins were tested for expression differences, of which 7 were differentially expressed carrying CS, heparan sulfate (HS) and/or keratan sulfate (KS) chains: versican core protein (CS), chondroitin sulfate proteoglycan 4 (CS), aggrecan core protein (CS/KS), testican-1 (HS/CS), syndecan-4 (HS/CS), collagen alpha-1(XVIII) chain (HS) and mimecan (KS). Expression differences of the 5 PGs carrying CS are shown on Fig. 4/B.

**Figure 4.**
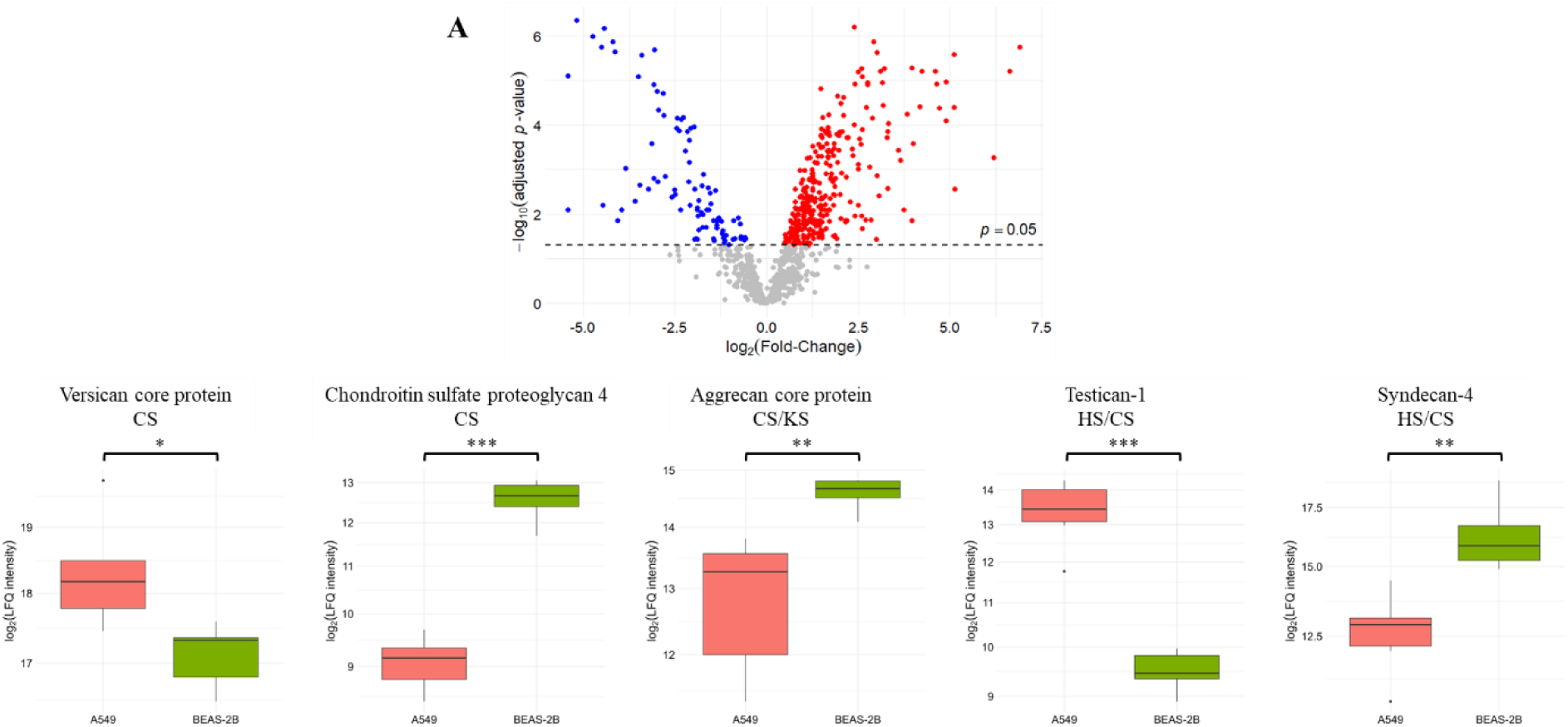
A. Volcano plot of all quantified proteins. Blue - significantly underexpressed in A549 sEVs, red - significantly overexpressed in A549 sEVs. **B.** Boxplots of differentially expressed CSPGs. (*p < 0.05, **p < 0.01, ***p < 0.001)

PCA was performed on the 945 proteins included in statistical analysis (Fig. 5/A), while hierarchical clustering was performed on the 408 differentially expressed proteins, and a heatmap was generated (Fig. 5/B). A549 and BEAS-2B sEVs separated completely based on principal component 1 (PC1 = 42.31%) in PCA and were clustered by group, indicating that the two sEV types display markedly different proteomic profiles.

**Figure 5.**
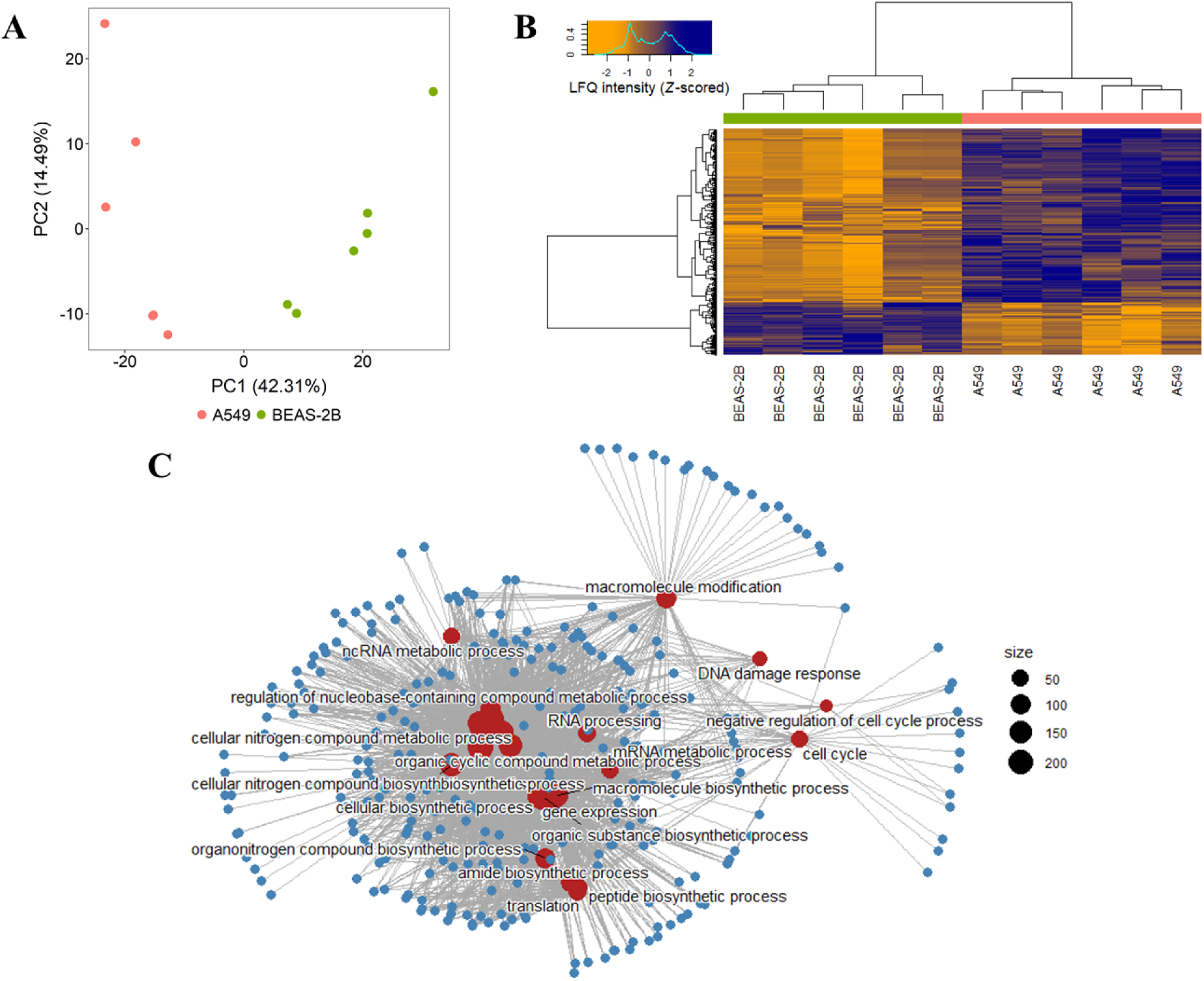
A. PCA analysis for all quantified proteins. **B.** Heatmap created after hierarchical clustering, generated for differentially expressed proteins. **C.** Gene-Concept Network based on the GSEA results for Gene Ontology Biological Process terms. Red nodes indicate processes, while blue nodes indicate genes.

Dysregulated processes were identified by GSEA and ranked based on their normalized enrichment scores (NES) and adjusted *p*-values (see Supplementary Table S-5). The top enriched process was negative regulation of cell cycle processes (NES = 1.75, p = 0.0321). Other highly ranked processes include cell cycle checkpoint signaling, cellular nitrogen compound metabolism, and regulation of DNA metabolic processes, all with NES values above 1.7. Additionally, processes related to nucleic acid and RNA metabolism, translation, and biosynthesis were significantly enriched. In contrast, negatively enriched processes include immune response, immune effector process and positive regulation of endocytosis, indicating downregulation. Fig. 5/C shows the linkages of genes and top GO biological process terms as a network.

### 3.3. *N*-glycoproteomics

In glycoproteomic analysis, 1731 *N*-glycopeptides – belonging to 513 peptides and 49 proteins – were quantified in the total of 18 samples (6 A549, 6 BEAS-2B, 3 F12 and 3 BEGM media). After grouping glycopeptides with the same glycan structure in the same position and excluding glycoforms found in less than 3 or detected in at least 2 media samples, a total of 301 glycoforms were included in statistical analysis, which are associated with 103 *N*-glycosylation sites of 46 proteins. 176 glycoforms (belonging to 71 glycosylation sites) were differentially represented between the two groups: 85 were overrepresented in A549 sEVs, while 91 were underrepresented in A549 sEVs. FCs and *p*-values are visualized in Fig. 6/A. For example, 13 glycoforms of versican CSPG core protein were differentially abundant. In position 1898, we observed the downregulation of 6 fucosylated complex *N*-glycans in A549, while 3 non-fucosylated tetra-antennary structures were upregulated. The full list of statistical results can be found in Supplementary Table S-6.

**Figure 6.**
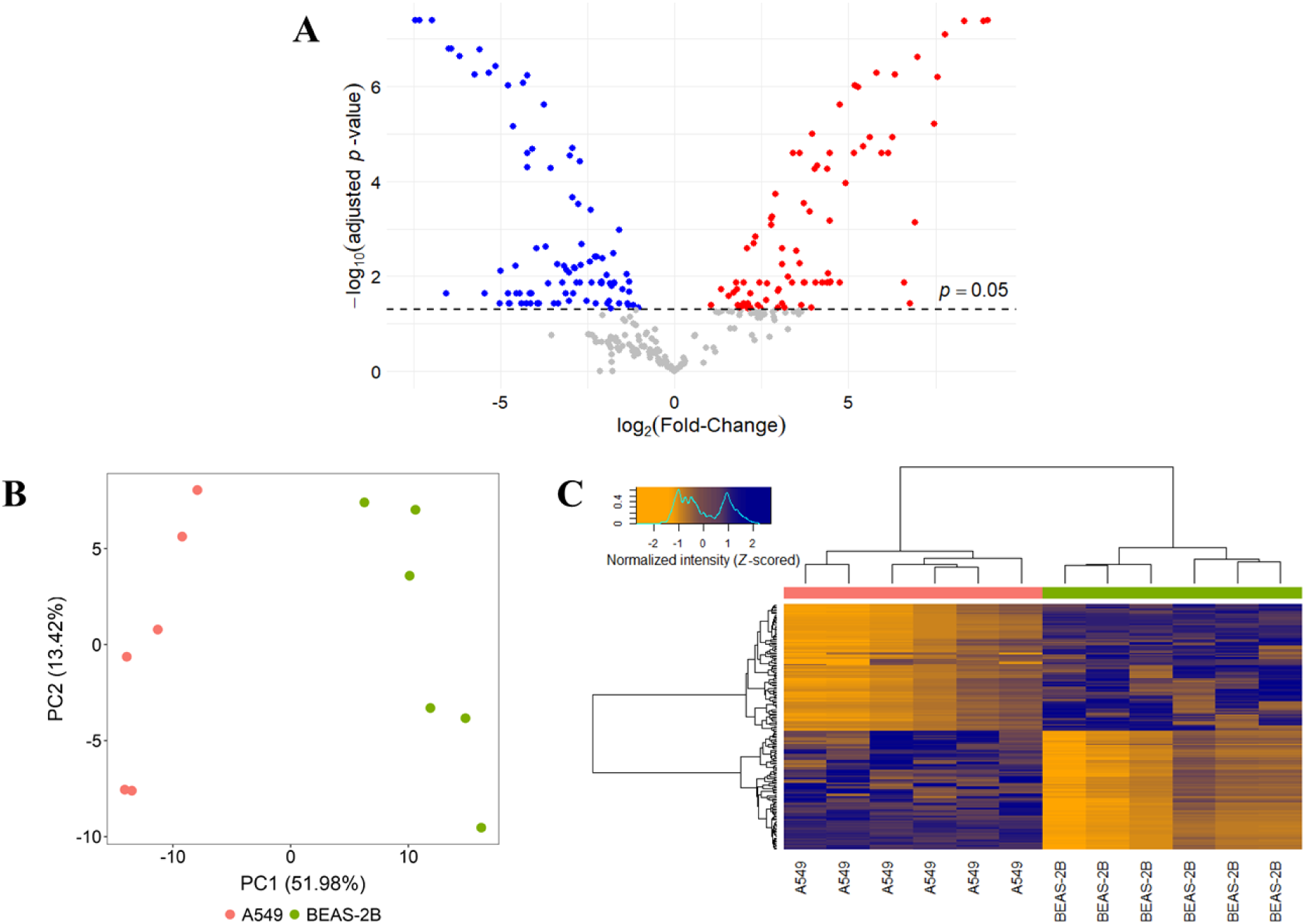
A. Volcano plot of all quantified glycoforms. Blue - significantly underrepresented in A549 sEVs, red - significantly overrepresented in A549 sEVs. **B.** PCA analysis for all quantified glycoforms. **C.** Heatmap created after hierarchical clustering, generated for differentially represented glycoforms.

PCA performed on all glycoforms subject to statistics shows that the two groups are completely separated based on PC1 (51.98%, Fig. 6/B), and heatmap analysis of differentially represented glycopeptides confirms that the two groups cluster separately (Fig. 6/C), indicating that sEVs derived from cancer and non-cancer cells have markedly different glycoproteomic profiles. To determine major glycosylation differences, glycans were characterized based on their type (complex, hybrid or oligomannose) and rates of fucosylation, galactosylation and sialylation (see Fig. 7, Supplementary Table S-7). In all samples, complex-type *N*-glycans were the most abundant (71% on average), while hybrid glycans were present in 26% on average and oligomannose glycans in 1.6%. However, in A549 sEVs, there was a 17% decrease in the abundance of complex glycans (*p* < 0.05) and a 37% decrease in the abundance of oligomannose glycans (non-significant), while the ratio of hybrid glycans increased (FC = 1.69, *p* < 0.05). A remarkable decrease (FC = 0.828, *p* < 0.05) was observed in fucosylation in case of A549, while galactosylation slightly increased (FC = 1.03, *p* < 0.05) and sialylation remained unchanged (FC = 1.01, non-significant).

**Figure 7.**
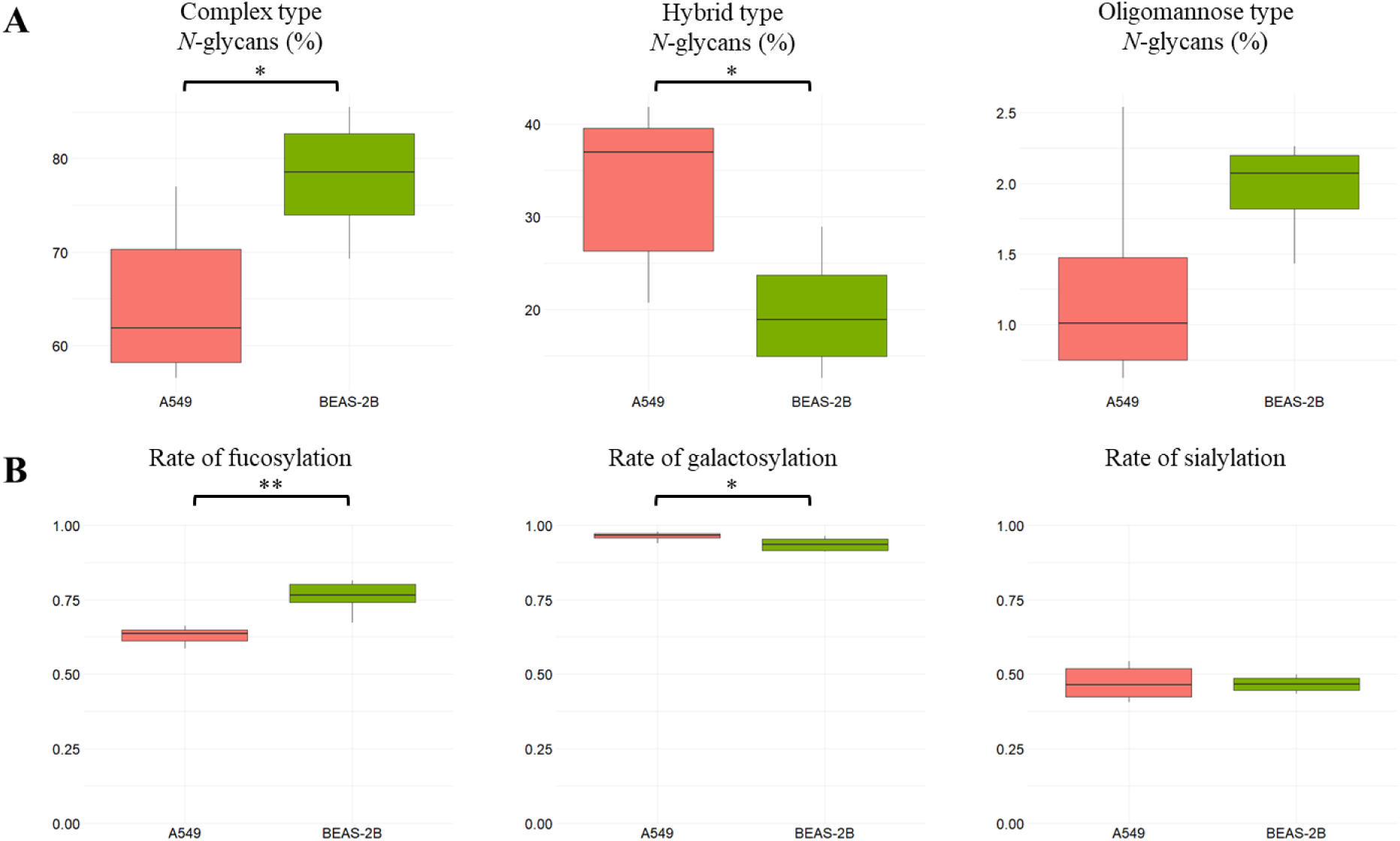
Boxplots of **A.** the ratio of different types of *N*-glycans (complex, hybrid and oligomannose). **B.** the rate of fucosylation, galactosylation and sialylation. (*p < 0.05, **p < 0.01, ***p < 0.001)

Changes in glycopeptide levels can result from changes in both protein levels and glycosylation characteristics. For example, galectin-3-binding protein (LG3BP) N551 F1H7N6S2 (indicating 1 fucose, 7 hexose, 6 *N*-acetylhexosamine and 2 sialic acid units) has a FC of 0.06, while FC of F1H5N4S1 at the same position is 5.0. The LG3BP protein also has a FC = 0.06 value, indicating that the former change is likely the result of protein level changes only, while the latter indicates glycan level changes. To eliminate differences in protein levels, the FC of the glycopeptide was normalized to the FC of the protein observed in proteomics. This allowed us to characterize a total of 206 glycoforms from 73 glycosylation sites of 26 proteins (listed in Supplementary Table S-8) that were analyzed in both glycopeptide analysis and in proteomics. Normalized GP FCs are visualized in Fig. 8, where small bubbles indicate that the observed glycopeptide change was caused by a change in protein amount, while larger bubbles indicate that a change in glycan structure occurred. On various laminin proteins (subunit alpha-2, LAMA2; subunit alpha-5, LAMA5; subunit gamma-1, LAMC1), mainly increased proportions of glycan structures were observed, e.g. in the case of LAMA2 N1810, 3 non-fucosylated structures were present in highly increased ratio. For mucin-5AC (MUC5A) and pappalysin-1 (PAPP1) proteins, a decrease in the rate of glycosylation was mainly observed. On some proteins, changes in both directions were detected, e.g. on versican (CSPG2) the ratio of glycan structures decreased at position N1898, while it increased at the other 3 positions analyzed.

**Figure 8.**
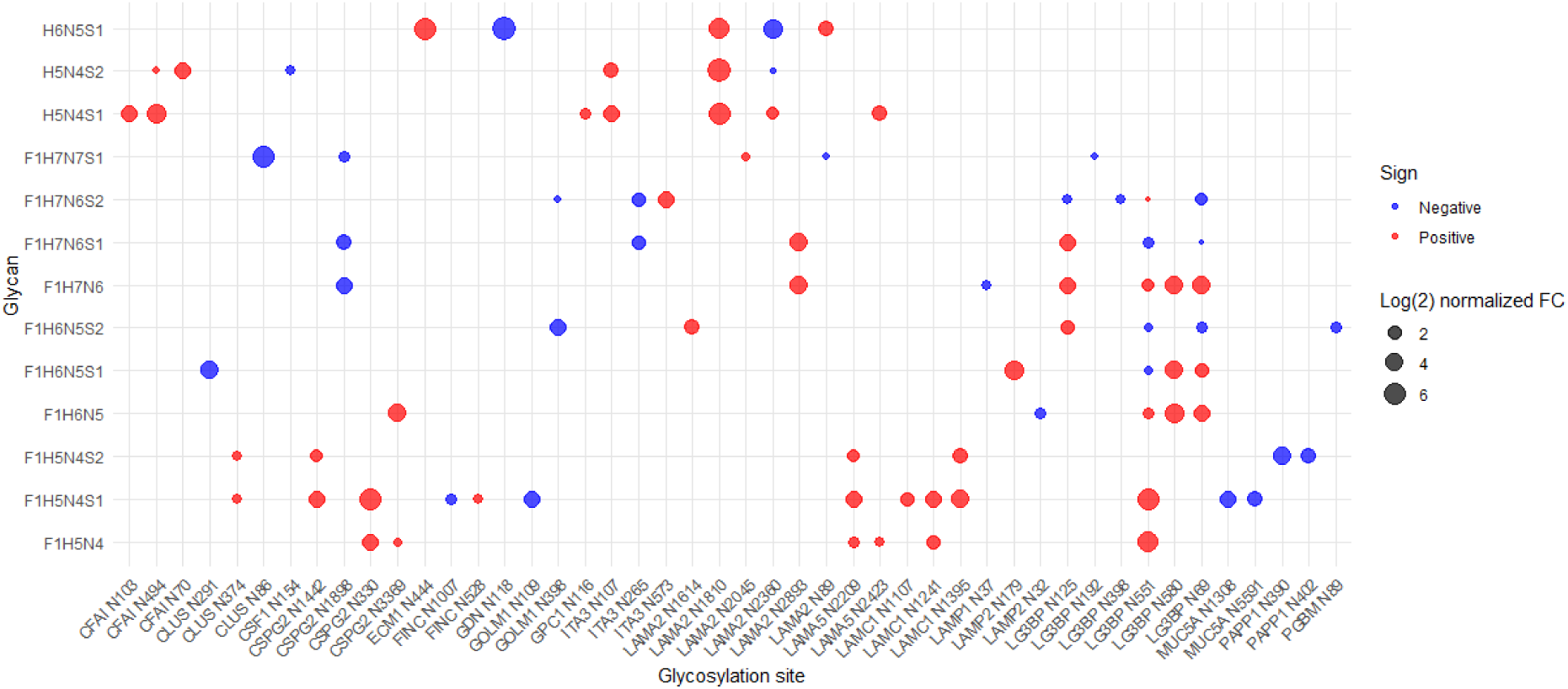
Bubble plot showing the normalized fold-change values of the glycoforms whose respective proteins were included in the proteomic statistical analysis. Red dots indicate an increase in the rate of glycosylation, blue dots indicate a decrease. Only glycans associated with at least 5 glycosylation sites are shown.

### 3.4. Chondroitin/Dermatan sulfate analysis

In CS/DS disaccharide analysis, no disaccharides were detected in either cell culture media, confirming that the CS/DS disaccharides measured in the samples were derived from sEVs. The disaccharide amounts in fmol calculated in the 12 EV sample are given in Supplementary Table S-9. In terms of the relative amounts of each component, the ratio of non-sulfated D0a0 (FC = 0.65) and monosulfated D0a6 (FC = 0.76) decreased in A549 sEVs compared to BEAS-2B sEVs, while that of monosulfated D0a4 (FC = 1.59) and disulfated D0a10 (FC = 1.23) increased (Fig. 9/A). Among the relative amounts of the 4 disaccharides, the change in the two monosulfated components was found to be statistically significant. These changes resulted in a slight increase in the average rate of CS/DS sulfation (Fig. 9/B, FC = 1.08, non-significant), while the 6S/4S ratio was significantly decreased in tumor sEVs (Fig. 9/C, FC = 0.48, significant), suggesting altered expression of sulfotransferase enzymes. On average, 3.4 times more CS/DS disaccharides were detected per sample in A549 sEVs than in BEAS-2B sEVs, illustrating that tumor samples contain more GAGs (Fig. 9/D). The statistical results for all calculated and derived quantities are presented in Supplementary Table S-10.

**Figure 9.**
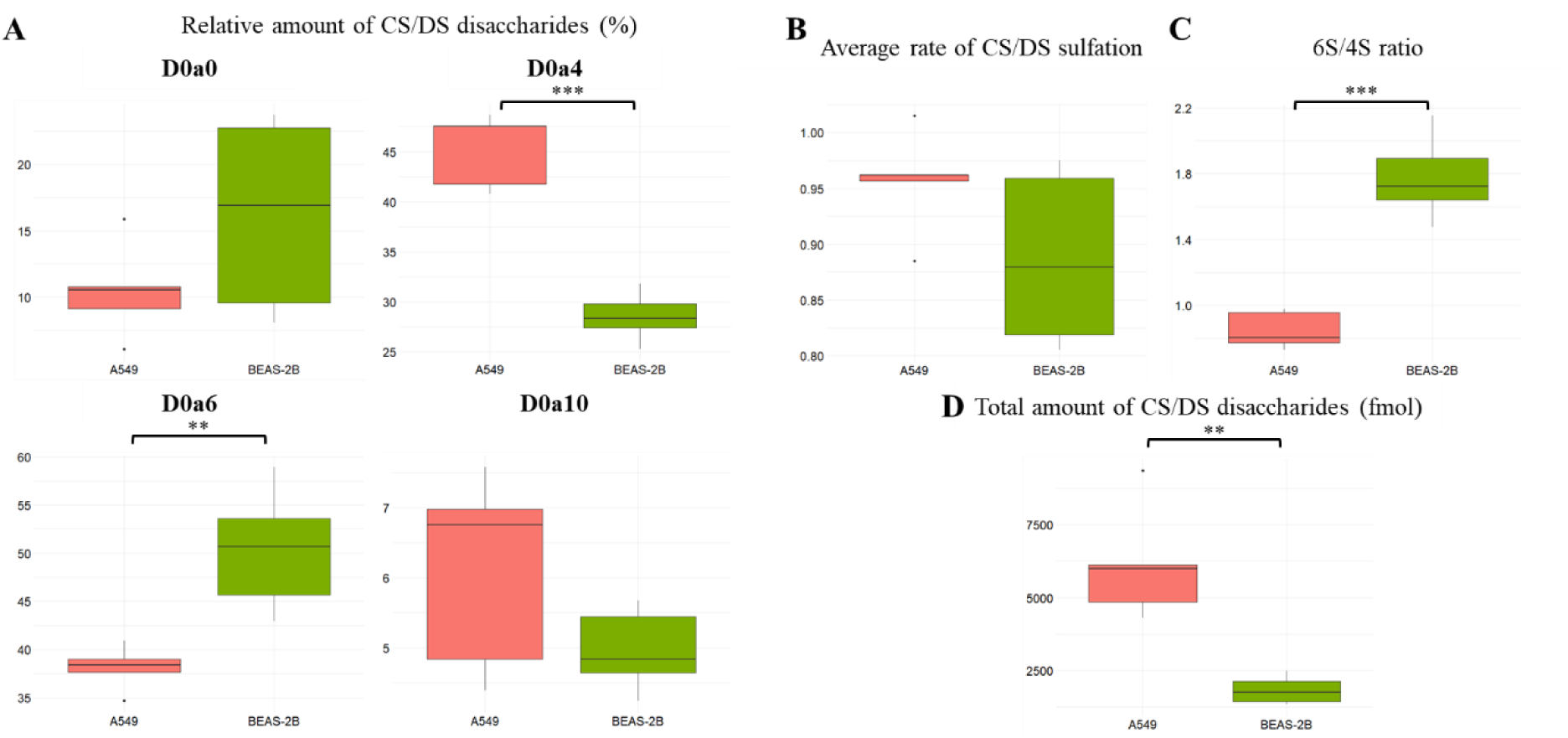
Boxplots of **A.** the relative amount of CS/DS disaccharides (%). **B.** the average rate of CS/DS sulfation. **C.** the 6S/4S ratio. **D.** the total amount of CS/DS disaccharides (fmol). (*p < 0.05, **p < 0.01, ***p < 0.001)

Next, CS/DS results were interpreted by hierarchical clustering and PCA for both relative (Fig. 10) and absolute (Supplementary Fig. S-2) amounts of each component. In the PCA analysis of relative CS/DS amounts, A549 and BEAS-2B sEVs were well distinguishable based on PC1 = 63.2% and PC2 = 29.04%, while in heatmap analysis, D0a4 and D0a10 disaccharides as well as D0a0 and D0a6 disaccharides clustered together, while samples clustered based on their classification.

**Figure 10.**
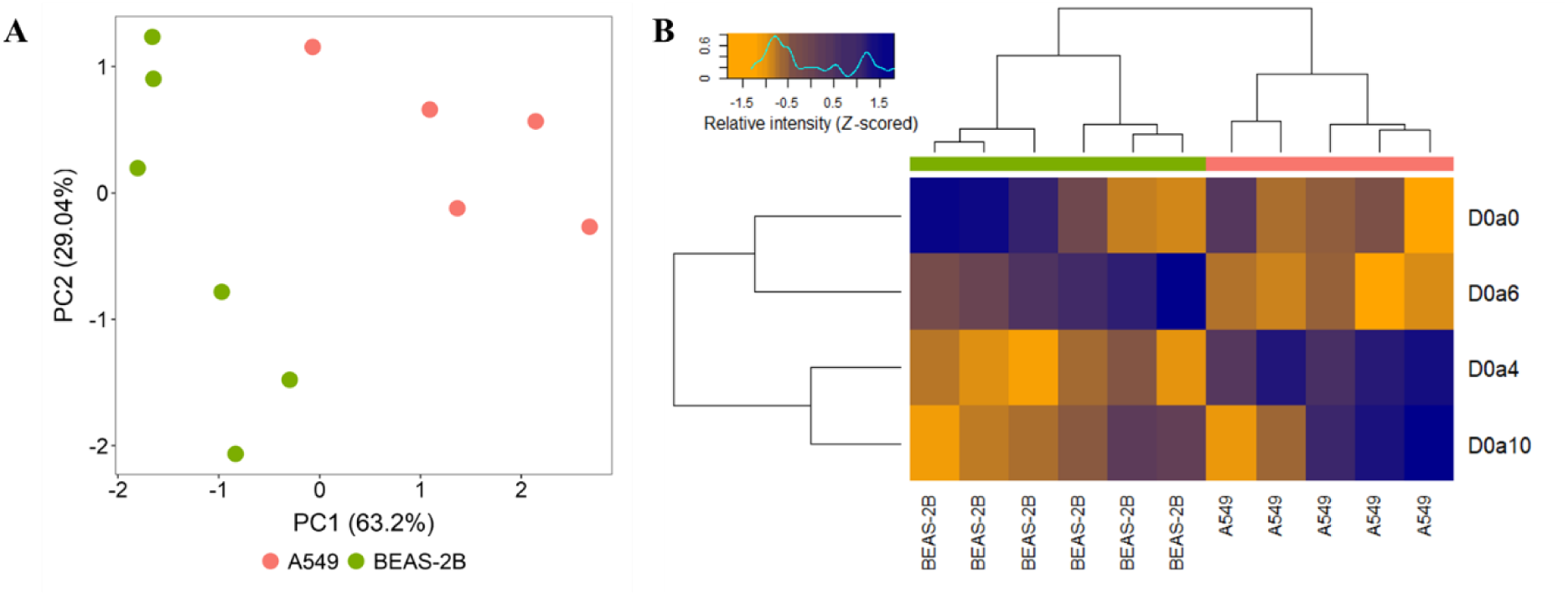
A. PCA analysis for the relative amounts of CS/DS disaccharides. **B.** Heatmap created after hierarchical clustering, generated for the relative amounts of CS/DS disaccharides.

## 4. Discussion

In the current study, we characterized the proteomic, *N*-glycoproteomic and CS/DS disaccharide profiles of sEVs derived from A549 and BEAS-2B cells. TEM and MRPS analysis confirmed the presence of particles <200 nm in size, and proteomic analysis confirmed the presence of common EV proteins, verifying the quality of sEVs.

### 4.1. Proteomics

The final proteomic dataset of 945 proteins allowed for robust statistical analysis, however, the presence of additional 583 proteins in at least two media samples highlights the challenge of background contamination, confirming the need for strict filtering criteria in EV proteomics. A high proportion of proteins (408 out of 945) was found to be differentially expressed, suggesting fundamental differences between the two sample types. The upregulation of several proteins found in the present study was previously observed in lung cancer tissue or plasma samples, e.g. fibronectin and proliferating cell nuclear antigen (PCNA). Fibronectin is an important constituent of the extracellular matrix (ECM) that supports cancer cell escape and cell migration leading to metastasis[53]. Increased fibronectin expression has been linked to poor prognosis in lung cancer, suggesting its role in tumor aggressiveness. Fibronectin levels have been previously shown to be significantly elevated in EVs from plasma of breast cancer patients compared to individuals without the disease, making it a promising marker for breast cancer detection[54]. PCNA largely reflects cell proliferation activity and is frequently overexpressed in lung tumors[55]. Beyond its role in proliferation, PCNA is involved in DNA synthesis, which may contribute to therapy resistance and tumor progression. Regarding the downregulated proteins observed, fibulin-2 is an ECM protein involved in cell adhesion and tissue organization that has been shown to be dysregulated in various cancers[56]. Analyzing lung cancer cell lines, fibulin-2 was downregulated in 9 out of 11 cell lines compared to normal bronchial epithelial cells, which was associated with DNA hypermethylation[57]. Complement C2, observed to be downregulated in lung cancer-derived sEVs, is a key component of the complement pathways, which help in immune surveillance and clearance of abnormal cells[58]. Therefore, reduced levels of C2 may impair complement activation and thereby promote tumor immune escape.

CSPG analysis identified alterations in both ECM PGs (versican, aggrecan, testican-1) and cell surface PGs (CSPG4, syndecan-4) in sEVs, which may reflect alterations in both tumor microenvironment and cell membrane-associated signaling of PGs[59]. CS, HS, and KS containing PGs were all affected, suggesting that all these GAG classes are worth investigating. The largest increase in cancer sEVs (FC = 14.3) was observed for testican-1, which has previously been shown to be upregulated in several cancer types, including lung cancer[60] and has been confirmed to be present in EVs[61]. High levels of this protein increase metastasis and reduce survival rates in cancer patients[62]. The largest decrease (FC = 0.083) was observed for syndecan-4, a protein that is upregulated or downregulated in a cancer type-dependent manner[63] and plays a key role in cell adhesion, migration, and signal transduction. Its downregulation has been linked to loss of adhesion, increased metastasis, and reduced responsiveness to extracellular signals, contributing to a more aggressive cancer phenotype[64]. GSEA revealed several dysregulated processes, that can be mostly associated with cell cycle regulation (e.g., negative regulation of cell cycle process, cell cycle checkpoint signaling), DNA repair (e.g., signal transduction in response to DNA damage, DNA damage response), metabolism (e.g., RNA metabolic process, nucleic acid metabolic process), protein synthesis (e.g., translation, peptide biosynthetic process), and immune response (e.g., immune response, immune effector process). Mostly upregulated processes were identified, while immune-related pathways showed negative enrichment. This pattern may partially reflect the higher number of upregulated proteins compared to downregulated ones in the dataset. The observed dysregulation of cell cycle-related processes is consistent with the well-known cancer hallmark of sustained proliferative signaling, where cancer cells bypass critical checkpoints to maintain uncontrolled proliferation[65]. Likewise, dysfunctional DNA repair mechanisms, another known feature of cancer, allow the accumulation of mutations, thereby leading to genetic diversity and the development of therapy resistance[66]. Cancer cells undergo significant metabolic reprogramming to meet the demands of rapid growth and survival, and are also characterized by elevated protein synthesis, which supports their aggressive growth and adaptation to the tumor microenvironment[67, 68]. Interestingly, we observed downregulation of immune-related processes in cancer EVs. This suggests immune evasion strategies employed by tumor cells, that allow them to evade host immune destruction while facilitating tumor progression[69].

### 4.2. *N*-glycoproteomics

In glycoproteomics, 301 glycoforms were analyzed, of which 176 were differentially represented between A549 and BEAS-2B sEVs. The observed profiles allowed for complete separation of the two groups in PCA, highlighting the distinct glycosylation patterns. This significant alteration in glycoform representation may reflect altered enzymatic activity of glycosyltransferases and glycosidases, which are frequently reported in tumorigenesis[28, 70]. For example, a previous study identified several glycosyltransferases that were associated with poor or good prognosis in cancer and may therefore be potential prognostic markers[71].

*N*-glycoproteomics and *N*-glycomics studies of EVs are very limited[72], with the scarce literature on EV glycoproteomics focusing mostly on urine[73] and blood plasma[74]. In lung cancer, a previous study compared *N*-glycomic patterns on sEVs from small cell lung cancer (SCLC) and non-small cell lung cancer (NSCLC) cells and found that the *N*-glycans of SCLC-sEVs are fairly heterogeneous, whereas NSCLC-sEVs contain primarily core-fucosylated, biantennary and triantennary *N*-glycans[75].

Interestingly, we observed an increased ratio of hybrid *N*-glycans and a decreased ratio of complex *N*-glycans in A549 sEVs. In cancer, it is well-documented that complex *N*-glycans, particularly those with highly branched structures, are commonly upregulated due to the overexpression of glycosyltransferases[70]. Our findings suggest incomplete glycan maturation in cancer EVs that may reflect altered glycosyltransferase expression or activity in A549 cells, e.g. reduced activity of *N*-acetylglucosaminyltransferases, responsible for branching of complex glycans, or an imbalance in Golgi processing enzymes could result in the accumulation of hybrid glycans[24].

Another remarkable observation was the significant decrease in fucosylation in A549 sEVs. Fucosylation plays critical roles in various biological processes, including cell signaling, adhesion, and immune modulation[76]. Core fucosyltransferase, responsible for core fucosylation, is frequently upregulated in various cancers[77], e.g. in lung cancer[78]. Based on our results, the reverse effect was detected in sEVs compared to glycosylation patterns commonly reported for cancer cells in the literature. Thus, further studies are needed to investigate the possible selective sorting of fucosylated glycans into EVs.

We characterized some glycoproteins that are highly glycosylated and several of their glycoforms were dysregulated in cancer sEVs, e.g. CSPG2, LG3BP and laminins.

CSPG2 is an ECM protein that plays a pivotal role in cell adhesion, migration, and tumor progression. Its G1 domain enhances cancer cell motility and reduces cell adhesion, while the G3 domain contributes to tumor invasiveness[79]. In our study, we identified several dysregulated CSPG2 glycoforms, including both up- and downregulations. At N1898 site, significant alterations were detected in the abundance of 9 glycoforms, of which 3 non-fucosylated, sialylated structures were overrepresented and 6 fucosylated forms were underrepresented in A549 sEVs. This change towards non-fucosylated structures indicates altered glycan sorting in A549 sEVs, potentially influencing their interactions within the tumor microenvironment and affecting processes like cell signaling and immune modulation.

LG3BP is a hyperglycosylated protein implicated in tumor progression, metastasis, and immune modulation[80]. It is enriched in cancer-associated EVs and is considered a promising candidate for targeted therapy in LG3BP-positive cancer[81]. A total of 23 differentially represented LG3BP glycoforms (at N69, N125, N192, N398 and N551) were detected in our study, all of which are underrepresented in A549 sEVs. However, compared to the LG3BP FC = 0.06 observed in proteomics, the majority of glycoforms indicate increased glycosylation rates.

Laminins are important ECM molecules involved in tumor angiogenesis, cell invasion and metastasis development, including the regulation of epithelial-mesenchymal transition and basement membrane remodeling[82]. Each of the laminin subunits LAMA2, LAMA5 and LAMC1 were found to possess dysregulated glycoforms. For example, LAMA2 at N1810 site showed the upregulation of bi- and triantennary structures, while hybrid glycans with H8N4 backbone were downregulated.

### 4.3. Chondroitin/Dermatan sulfate analysis

CS/DS disaccharide analysis revealed the increased amount of CS/DS chains in A549 sEVs compared to BEAS-2B sEVs (FC = 3.4). Increased amount of CS/DS in tumors is commonly observed in several types of cancer, e.g. in liver[34], prostate [33] and lung[83] cancer tissues. The average rate of CS/DS sulfation slightly increased (FC = 1.1), and we observed a marked difference in the 6S/4S ratio (FC = 0.48).

MS based research on GAG analysis of EVs is scarce. So far, only a specific GC-MS based technique was applied, which breaks all glyco-polimers, including GAGs, into their constituent saccharide units. Using this method, it was demonstrated that EVs derived from melanoma cells with or without brain metastasis contain different amounts of hyaluronan (HA)[84], and that EVs derived from plasma have different glycan profiles from whole plasma and are enriched in CS, DS and KS GAGs[85].

Thus, there is no literature available on the analysis of EV CS/DS disaccharides, but CS/DS GAG analysis of lung tissue samples has been previously performed. In a comprehensive study of CS/DS, HS, and HA, the cancer tissue samples contained over twice as much CS/DS as did the normal tissue samples and the 6S/4S ratio greatly increased (FC = 2.0), while the amount of HS and HA were not significantly different[32]. Examining the CS/DS characteristics of tumor and adjacent normal regions from patients with different types of lung cancer, the total amount of CS/DS disaccharides was higher (FC = 2.2) in tumor than in adjacent normal regions; the relative amount of D0a0 decreased (FC = 0.75), while the amount of monosulfated components (D0a4, FC = 4.1 and D0a6, FC = 2.3) increased in tumor[83]. This resulted in an increase in the average CS/DS sulfation (FC = 2.4) and a decrease in the 6S/4S ratio (FC = 0.83). In ALK rearranged lung adenocarcinoma tissues, total CS/DS increased by 2.5-4.4-fold depending on the sample type, while average sulfation increased by 4.0-4.7-fold and 6S/4S sulfation showed variability but was mostly increasing[86]. Compared to these findings in literature, there is a large increase in CS/DS amount in lung cancer sEVs, similar to tissues, but we did not observe a marked increase in the rate of sulfation, and the 6S/4S ratio in our study was significantly decreased in cancer, whereas previous studies have shown varying trends in tissues.

Changes in the sulfation pattern, as observed in our study, play an important role in regulating cell signaling pathways and are therefore strongly linked to cancer progression and metastasis[87]. The decreased 6S/4S ratio suggests the overexpression of chondroitin 4-*O*-sulfotransferase enzymes, mainly carbohydrate sulfotransferase 11. The overexpression of this gene has been associated with unfavorable prognosis in the case of liver[88], pancreatic[89], and lung[90] cancer.

Oncofetal CS (ofCS) is a unique GAG structure that has been associated with both fetal development and cancer[91]. For example, high levels of ofCS in non-small cell lung cancer tissues have been linked to poor patient survival[92]. OfCS can be detected using the recombinant VAR2CSA protein, which specifically recognizes this altered GAG structure. One of its defining features is an increase in 4-*O*-sulfation[93]. Based on our observations, it is possible that tumor-derived sEVs carry ofCS, potentially contributing to the high levels of 4-*O*-sulfation detected in our study. However, there is currently no evidence to support this hypothesis.

## Conclusions

Our study provides novel insights into the proteomic, *N*-glycoproteomic and CS/DS GAG profiles of sEVs derived from A549 lung adenocarcinoma and BEAS-2B non-tumorigenic cell lines. While proteomic characterizations of sEVs are widely available in literature, glycosylation and GAG profiles remain underexplored.

The findings show that all three analyzed profiles largely reflect sEV origin, as hierarchical clustering and PCA consistently distinguish cancer-derived sEVs from non-cancerous ones. Proteomics revealed significant dysregulation of 5 CSPGs, such as testican-1 and syndecan-4, which are involved in processes such as cell adhesion, migration and tumor progression.

In *N*-glycoproteomic analysis, we observed a decrease in the rate of fucosylation and complex glycans in A549 sEVs. This pattern contrasts with the changes observed at the cellular level in general, suggesting selective glycan selection in sEVs.

CS/DS analysis revealed a 3.4-fold increase in total CS/DS disaccharide content in cancer sEVs, accompanied by an altered 6S/4S ratio. This altered sulfation pattern may modulate the interactions of sEV with receptors in the tumor microenvironment.

To conclude, our study highlights the significant proteomic and glycosylation differences between cancer and non-cancer cell derived sEVs, emphasizing the potential of these molecular signatures as diagnostic markers. Future studies on sEVs from patient biofluids may validate these findings and pave the way for their clinical application in liquid biopsy-based cancer diagnostics.

## Author contributions

Mirjam Balbisi: Data Curation, Formal Analysis, Investigation, Methodology, Visualization, Writing – Original Draft Preparation

Tamás Langó: Investigation, Methodology, Writing – Review & Editing

Virág Nikolett Horváth: Investigation, Formal Analysis, Writing – Review & Editing

Domonkos Pál: Investigation, Writing – Review & Editing

Gitta Schlosser: Resources, Writing – Review & Editing

Gábor Kecskeméti: Investigation, Writing – Review & Editing

Zoltán Szabó: Resources, Writing – Review & Editing

Kinga Ilyés: Investigation, Writing – Review & Editing

Nikolett Nagy: Investigation, Writing – Review & Editing

Otília Tóth: Investigation, Writing – Review & Editing

Tamás Visnovitz: Investigation, Resources, Writing – Review & Editing

Zoltán Varga: Resources, Writing – Review & Editing

Beáta G. Vértessy: Resources, Writing – Review & Editing

Lilla Turiák: Conceptualization, Funding Acquisition, Methodology, Project Administration, Supervision, Writing – Review & Editing

## Acknowledgments

Proteomic samples were run at the ISTA LSF Mass Spectrometry Facility, part of the Scientific Service Units of ISTA. The project was supported by the Lendület (Momentum) Program of the Hungarian Academy of Sciences (HAS, MTA), the National Research, Development and Innovation Fund of Hungary (K135231, K146890, 2022-1.2.2-TÉT-IPARI-UZ-2022-00003), the TKP2021-EGA-02, the TKP2021-EGA-31 and the 2020-1-1-2-PIACI-KFI_2020-00021 grants, implemented with support provided by the Ministry for Innovation and Technology of Hungary from the National Research, Development and Innovation Fund, and the ICGEB Research Grants Programme 2023 (CRP/HUN23-02). Project no. SNN 148580 has been implemented with the support provided by the Ministry for Innovation and Technology of Hungary from the National Research, Development and Innovation Fund, financed under the SNN_24 funding scheme. MB and DP were supported by the Semmelweis 250+ Excellence PhD Scholarship. TV was supported by VEKOP-2.3.3-15-2017-00016, RRF-2.3.121-2022-00003, TKP2021-EGA-23 and the János Bolyai Research Fellowship of the Hungarian Academy of Sciences. OT was supported by the Doctoral Excellence Fellowship Programme (DCEP) funded by the National Research, Development and Innovation Fund of the Ministry of Culture and Innovation and the Budapest University of Technology and Economics.

## Data availability statement

The data of the proteomic and N-glycoproteomic measurements are available in the MassIVE repository under the https://doi.org/doi:10.25345/C51834F3N link and can be downloaded via FTP (ftp://massive.ucsd.edu/v09/MSV000097305/). The GAG-omics data presented in this study have been deposited in the GlycoPOST database[94] under the accession number of GPST000562.

