## Supplementary figures for "A549 tumorigenic and BEAS-2B non-tumorigenic cell line derived small extracellular vesicles show distinct proteomic, *N*-glycoproteomic and chondroitin/dermatan sulfate profiles"

### List of supplementary figures

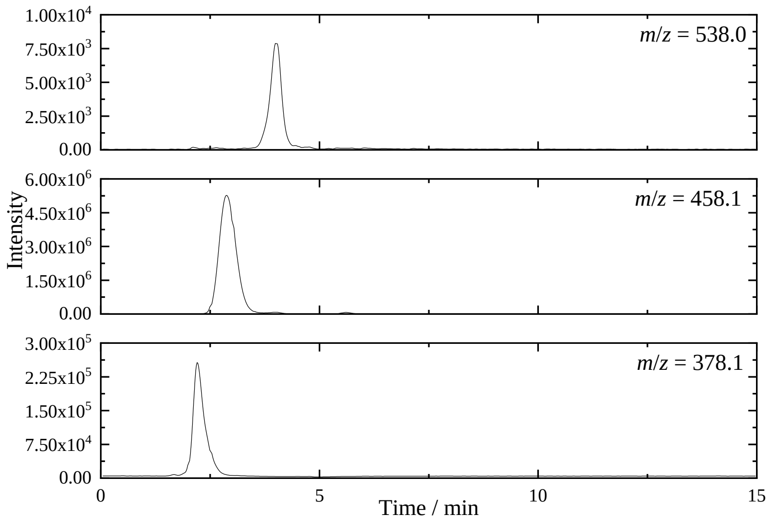


**Figure S-1.** Extracted ion chromatograms of CS/DS disaccharides.

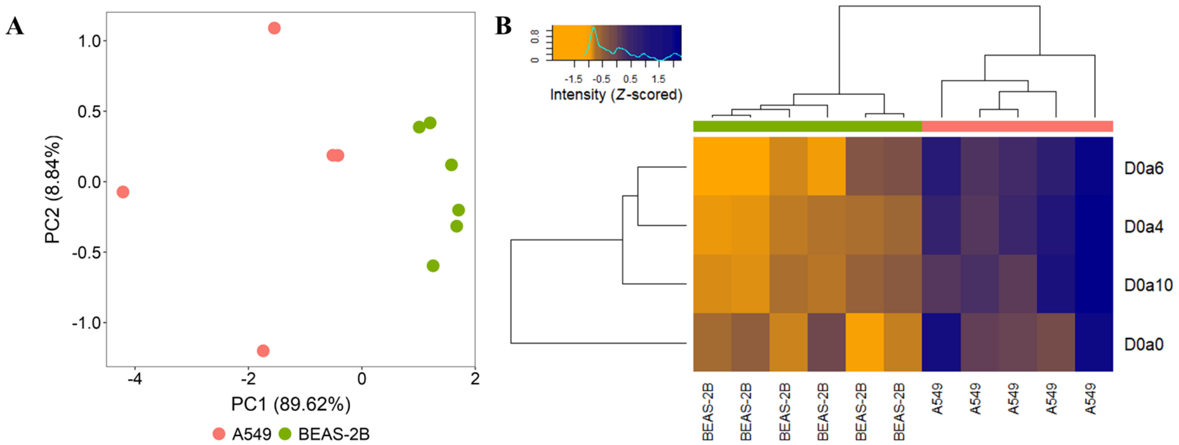


**Figure S-2. A.** PCA analysis for the total amounts of CS/DS disaccharides. **B.** Heatmap created after hierarchical clustering, generated for the total amounts of CS/DS disaccharides.

**List of supplementary tables** (in separate excel sheets)

**Table S-1.** Passage numbers of cells used for EV isolations.

**Table S-2.** Software search settings.

**Table S-3.** Glycan database used for GlycReSoft analysis.

**Table S-4.** List of proteins included in statistical analysis.

**Table S-5.** List of altered biological processes according to GSEA analysis.

**Table S-6.** List of glycoforms included in statistical analysis.

**Table S-7.** Statistical results for *N*-glycoproteomics metrics.

**Table S-8.** Glycoform intensities normalized by protein amounts.

**Table S-9.** Total amount of each CS/DS disaccharide per sample [fmol].

**Table S-10.** Statistical results for CS/DS disaccharide metrics.
